## Supplemental Material for "DeepAtlas: a tool for effective manifold learning"

Serena J. Hughes *et al.*

**This PDF file includes:**

Figs. S1 to S16

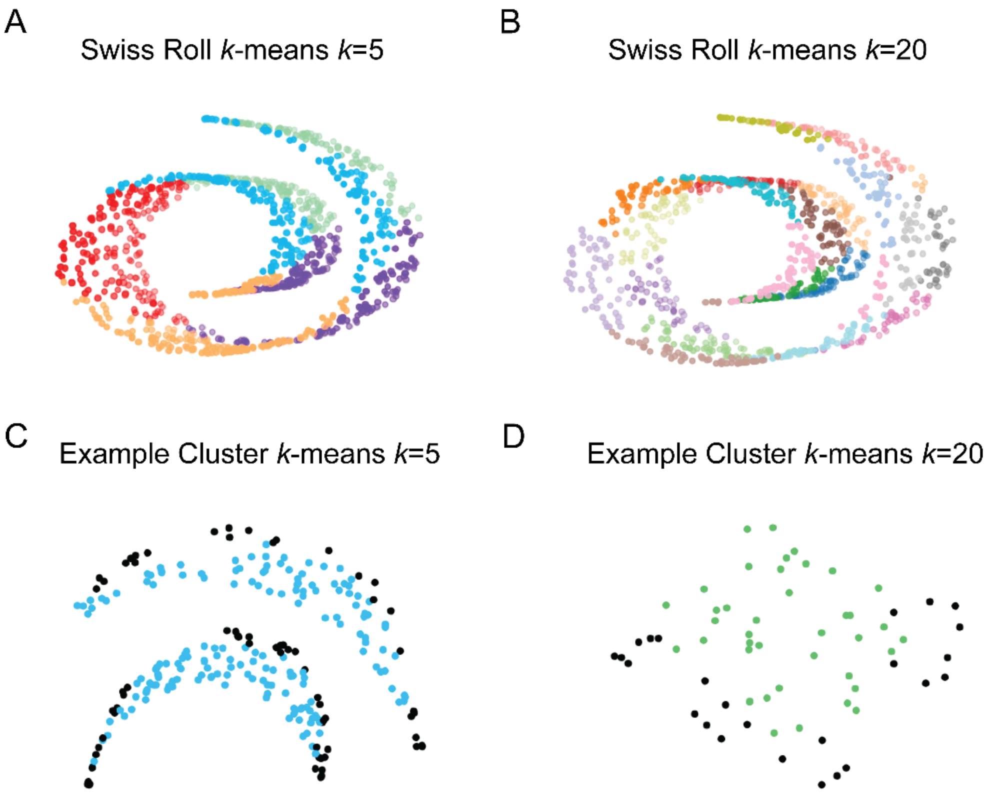


**Fig. S1.**
A) Swiss roll dataset in 3D with points colored based on $k$-means clustering using $k = 5$. B) Swiss roll dataset in 3D with points colored based on $k$-means clustering using $k = 20$. C) Example of one cluster from the Swiss roll data shown in S1A where points that are not nearby in the original dataset have been clustered together. D) Example cluster from the Swiss roll data shown in S1B. While some clusters still contain points that aren't neighbors in the 3D data, using more clusters reduces the occurrence.

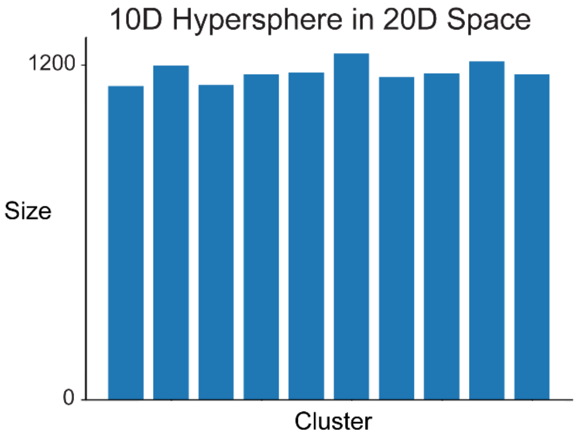


**Fig. S2.**
Cluster composition of the $k$-means clusters of the 10-dimensional hypersphere data embedded in 20-dimensional space, using $k = 10$.

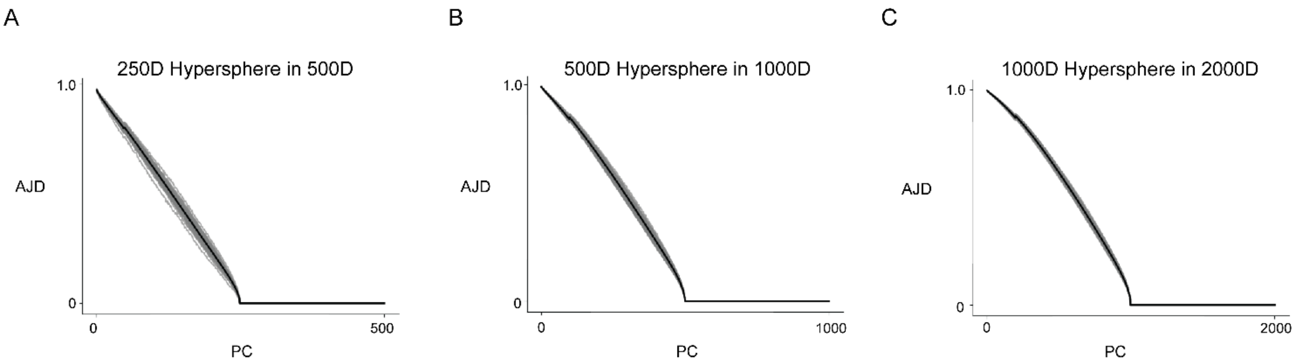


**Fig. S3.** PC vs. AJD for higher dimensional hyperspheres. A) 250-dimensional hypersphere of 5,000 points embedded in 500-dimensional space. B) 500-dimensional hypersphere of 10,000 points in 1000D space. C) 1000-dimensional hypersphere of 20,000 points in 2000-dimensional space.

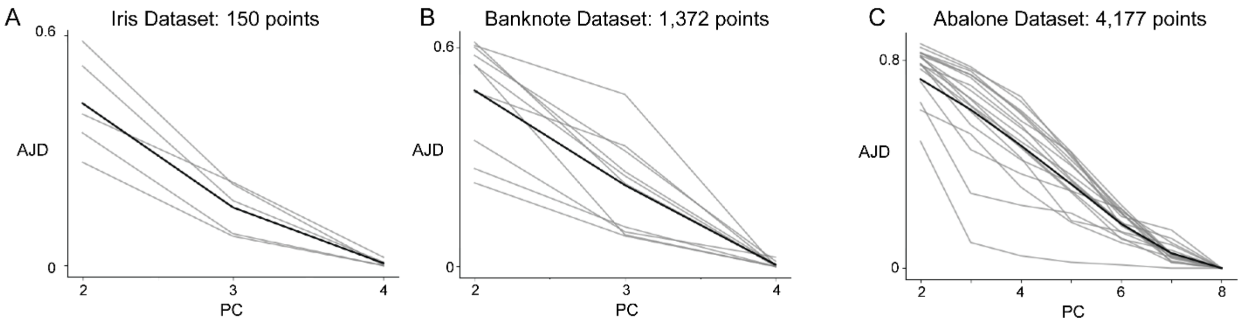


**Fig. S4.**
PC vs. AJD plots of other standard test machine learning datasets which do not indicate a lower dimensional manifold. A) Iris dataset with 4 dimensions and 150 points. B) Banknote dataset with 4 dimensions and 1,372 points. C) Abalone dataset with 8 dimensions and 4,177 points.

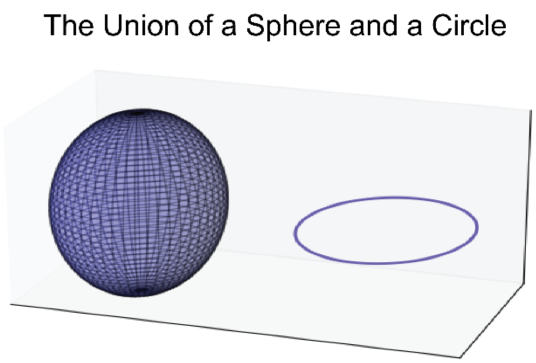


**Fig. S5.**
One dataset consisting of both a sphere and a circle, each of which is its own manifold.

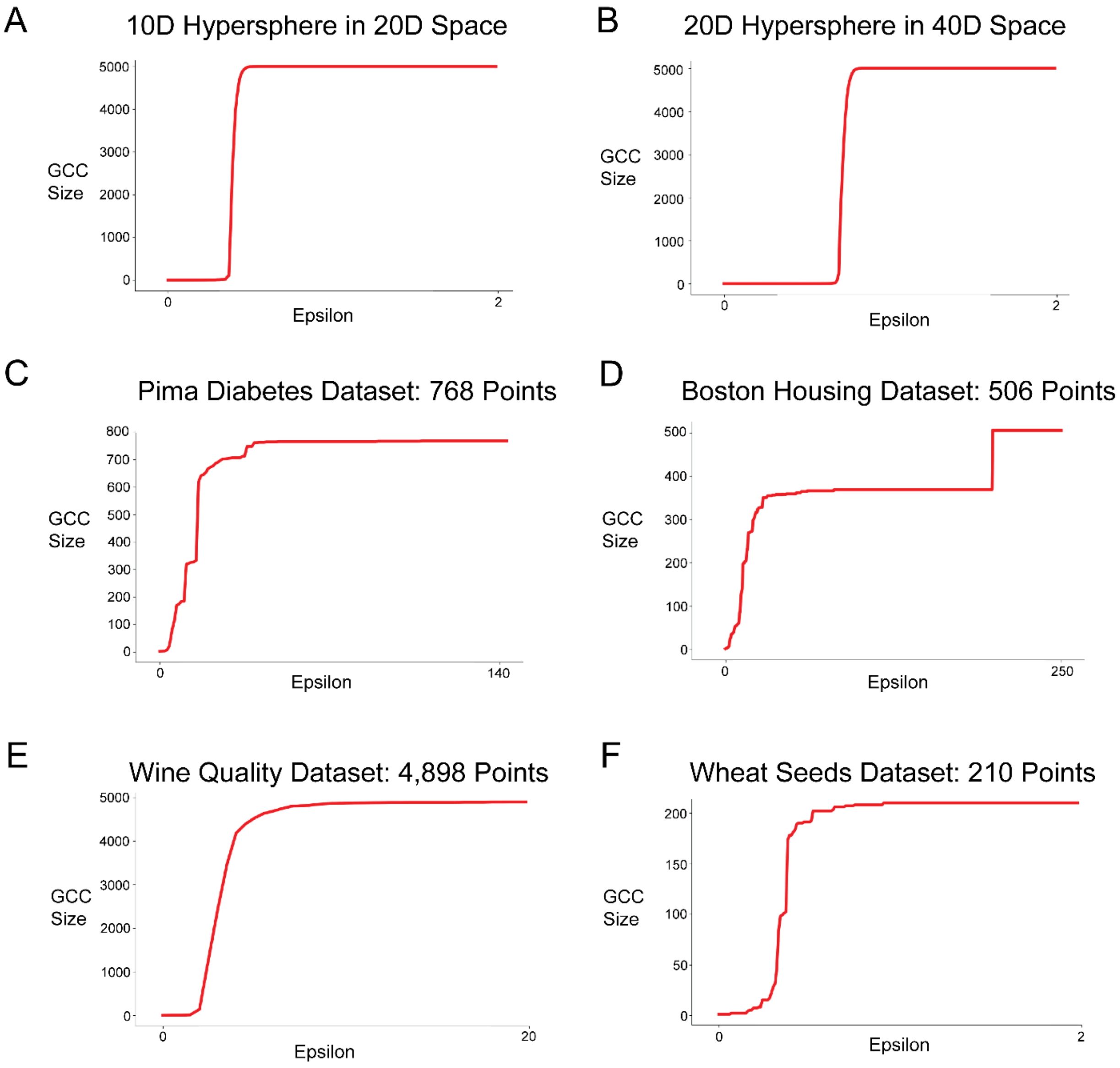


**Fig. S6.**
Epsilon plotted against the size of the giant connected component for each dataset. Smooth curves indicate the data is composed of one connected component while step behavior shows multiple components that are being joined together.

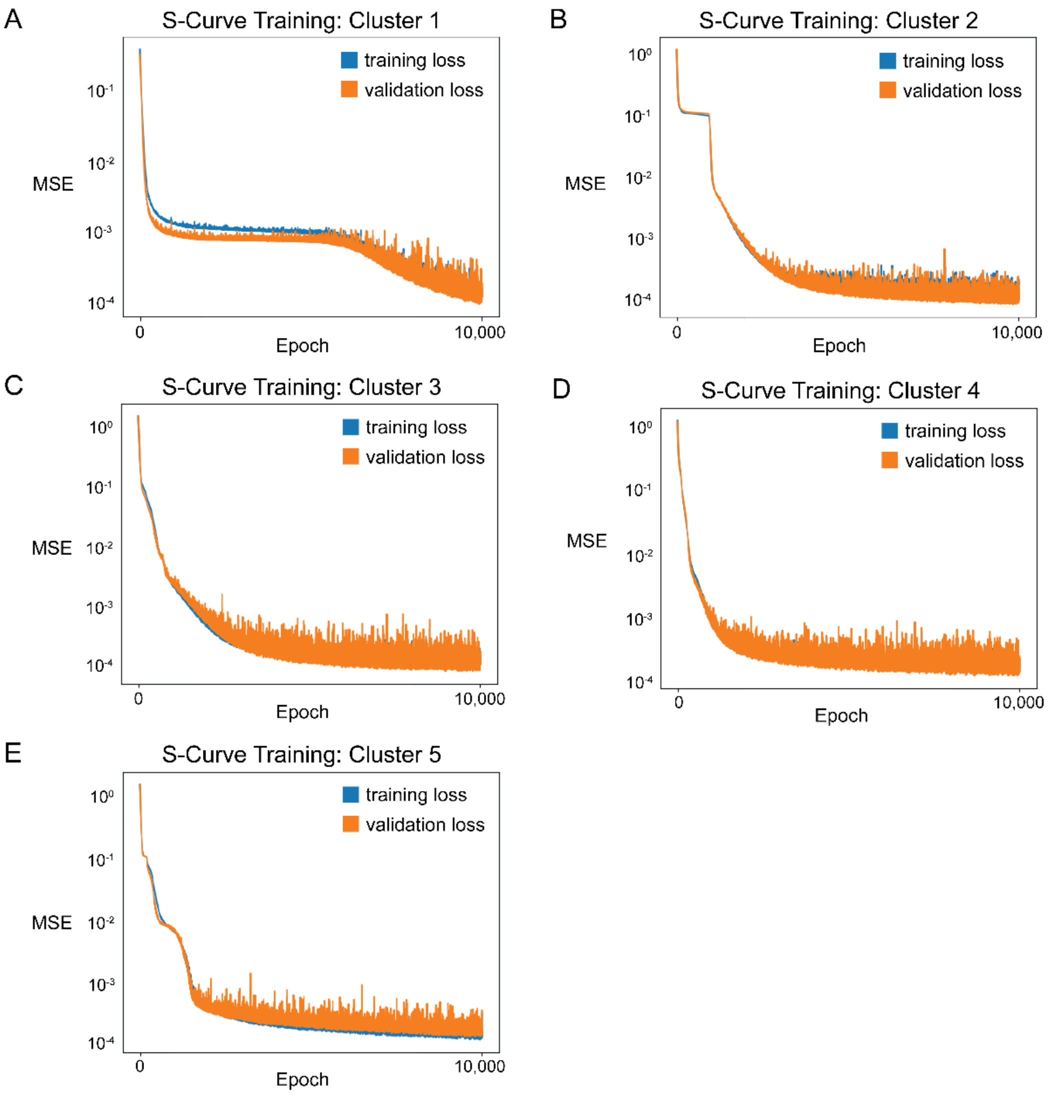


**Fig. S7.**
Training results of DeepAtlas' neural network on each of the S-curve clusters, generated using $k$-means with $k = 5$.

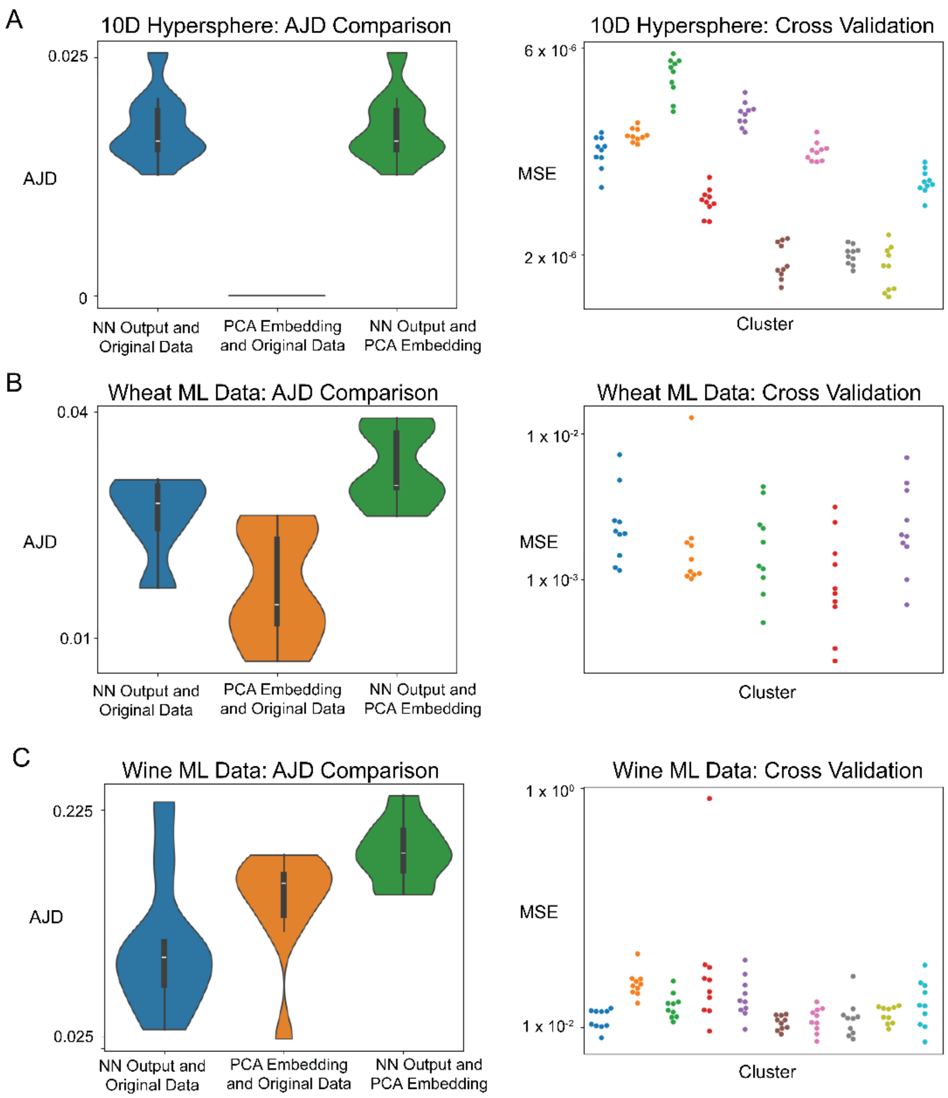


**Fig. S8.**
AJD comparison and 10-fold cross validation results for DeepAtlas applied to other datasets which indicated a lower dimensional manifold. A) 10D Hypersphere embedded into 20D space, with a lower dimension at 10D. B) Wheat machine learning test data with a lower dimension at 4D. C) Wine machine learning test data with a lower dimension at 10D.

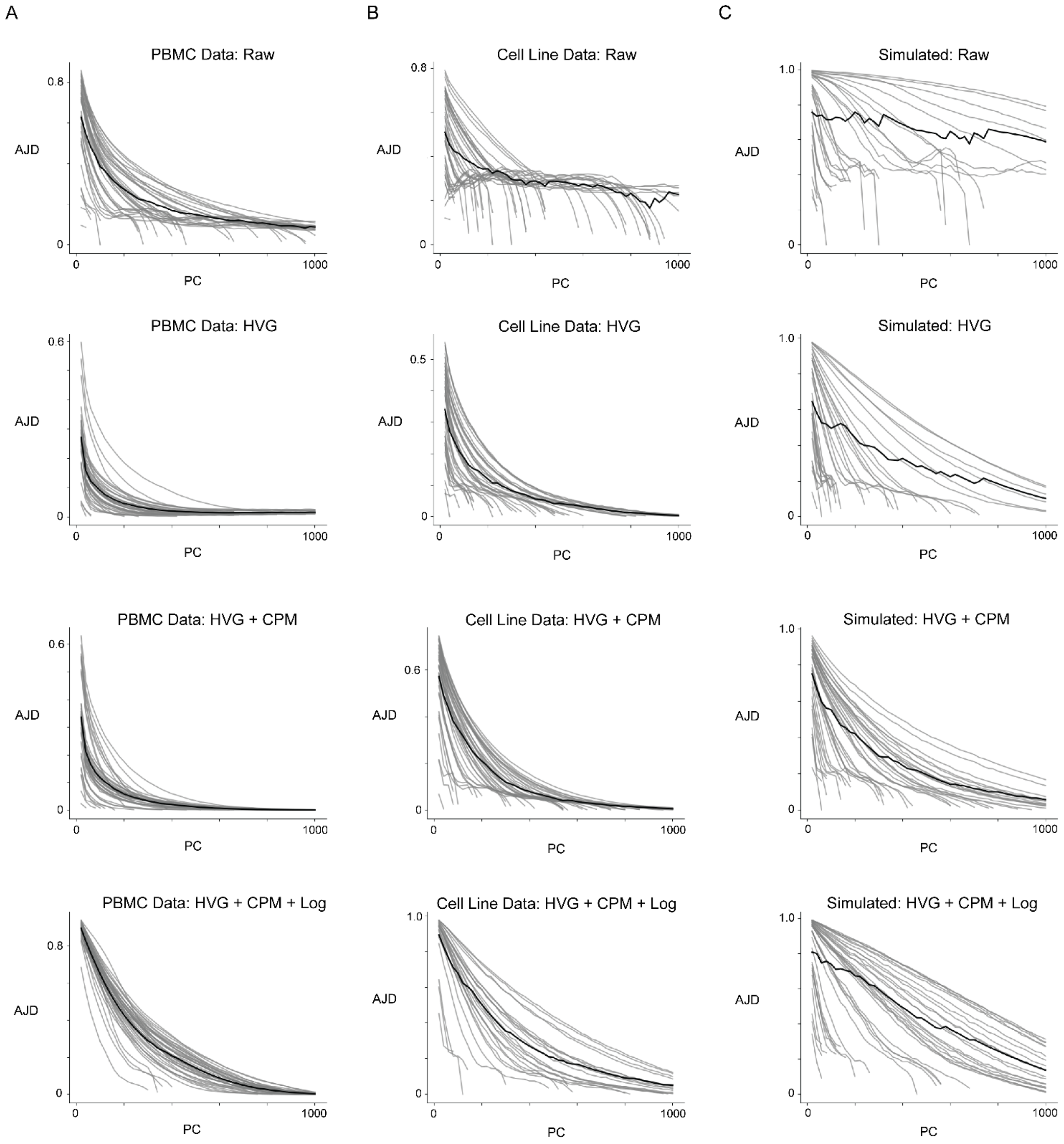


**Fig. S9.**
PC vs. AJD plots for additional single cell RNA Sequencing datasets explored, across all preprocessing steps. A) The full PBMC data. B) Cell Line Data. C) Simulated data generated using sc-Design, based on the cell line data.

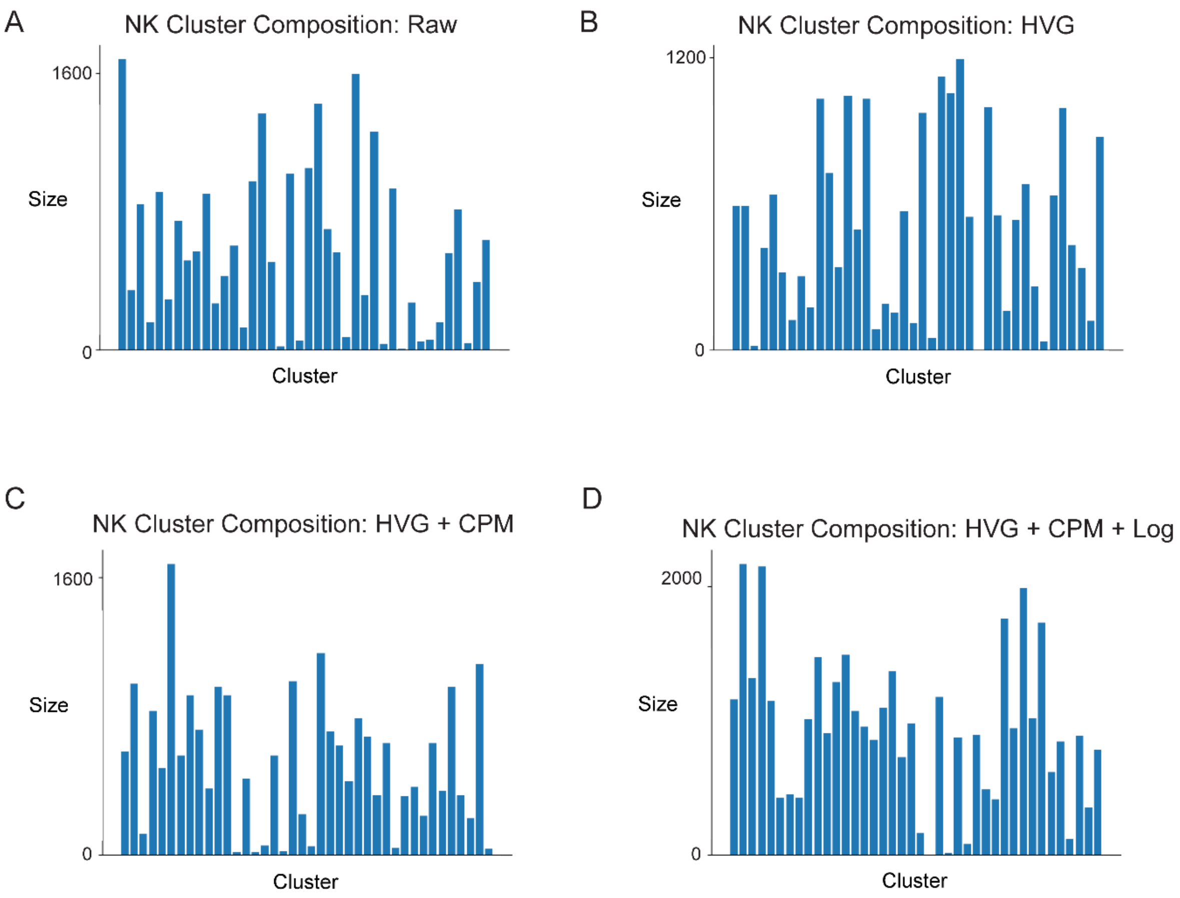


**Fig. S10.**
NK cell scRNA-seq data cluster sizes across preprocessing stages. The trend of inconsistent sizing occurs in each.

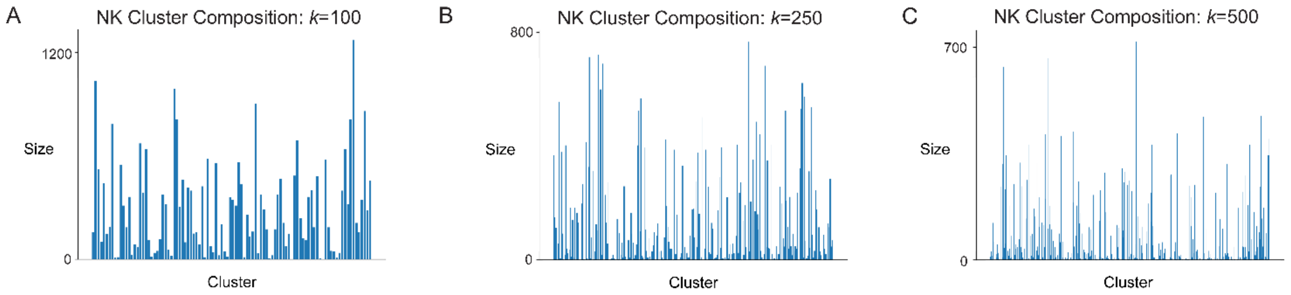


**Fig. S11.**
Raw NK cell scRNA-seq data cluster sizes when using different values of $k$ for $k$-means. See S10 for $k = 40$.

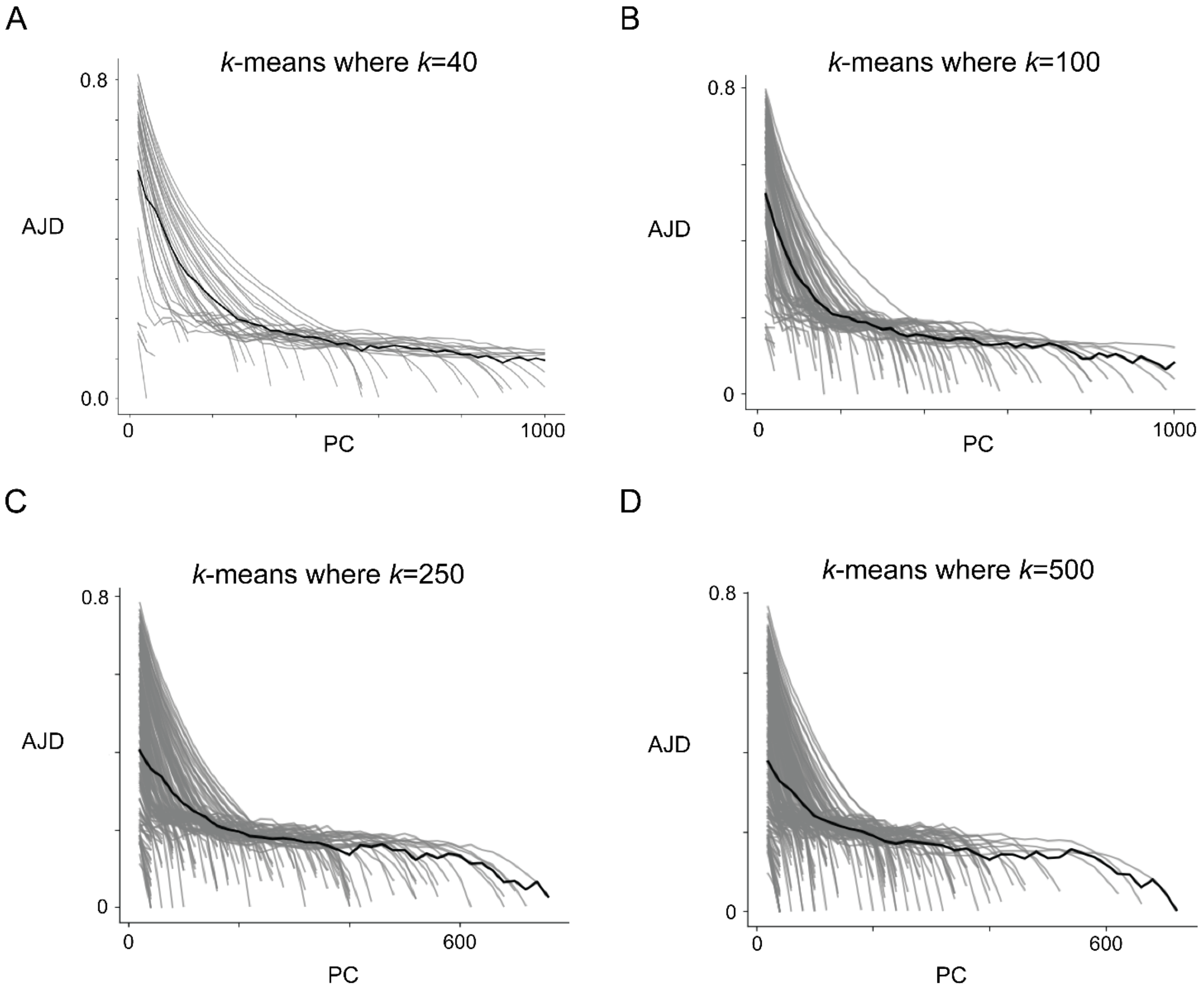


**Fig. S12.**
PC vs. AJD plots for the raw NK cell scRNA-seq data when using different values of $k$ for the $k$-means. A) $k = 40$, as shown in the main text. B) $k = 100$. C) $k = 250$. D) $k = 500$.

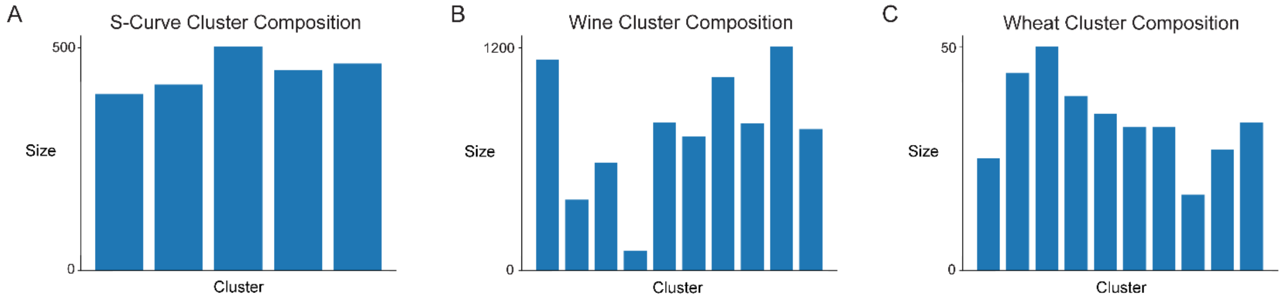


**Fig. S13.**
Cluster sizes when $k$-means is applied to other datasets which did indicate a lower dimensional manifold. A) S-Curve data with $k = 5$. B) Wine machine learning data with $k = 10$. C) Wheat machine learning data with $k = 10$.

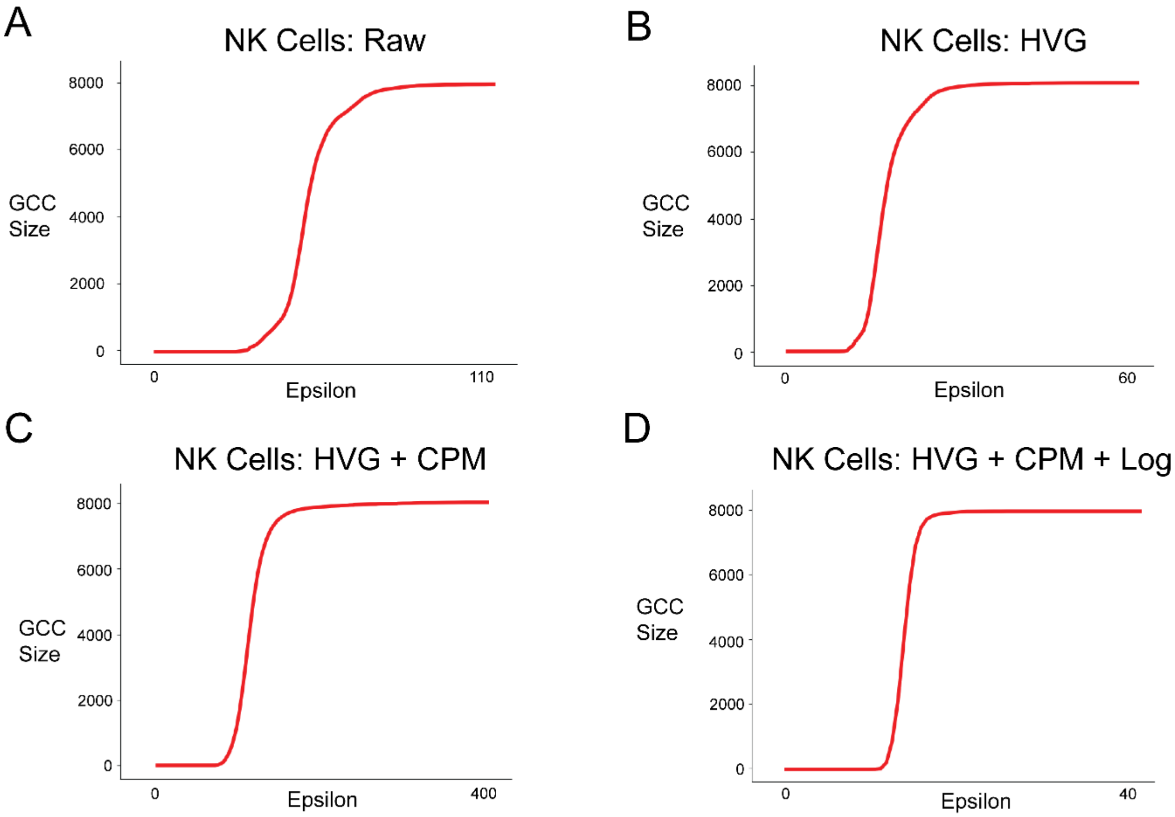


**Fig. S14.**
Epsilon plotted against the giant connected component size for the NK cell scRNA-seq data at each preprocessing stage.

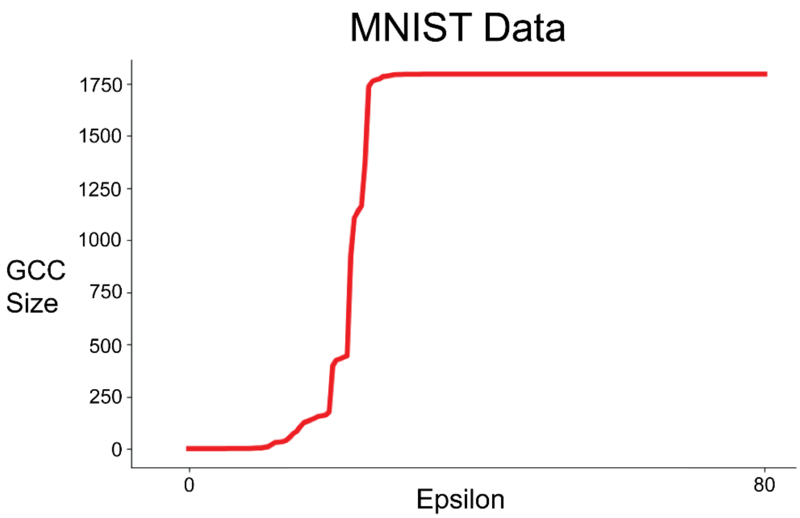


**Fig. S15.**
Epsilon plotted against the giant connected component size for the MNIST handwritten digits data.

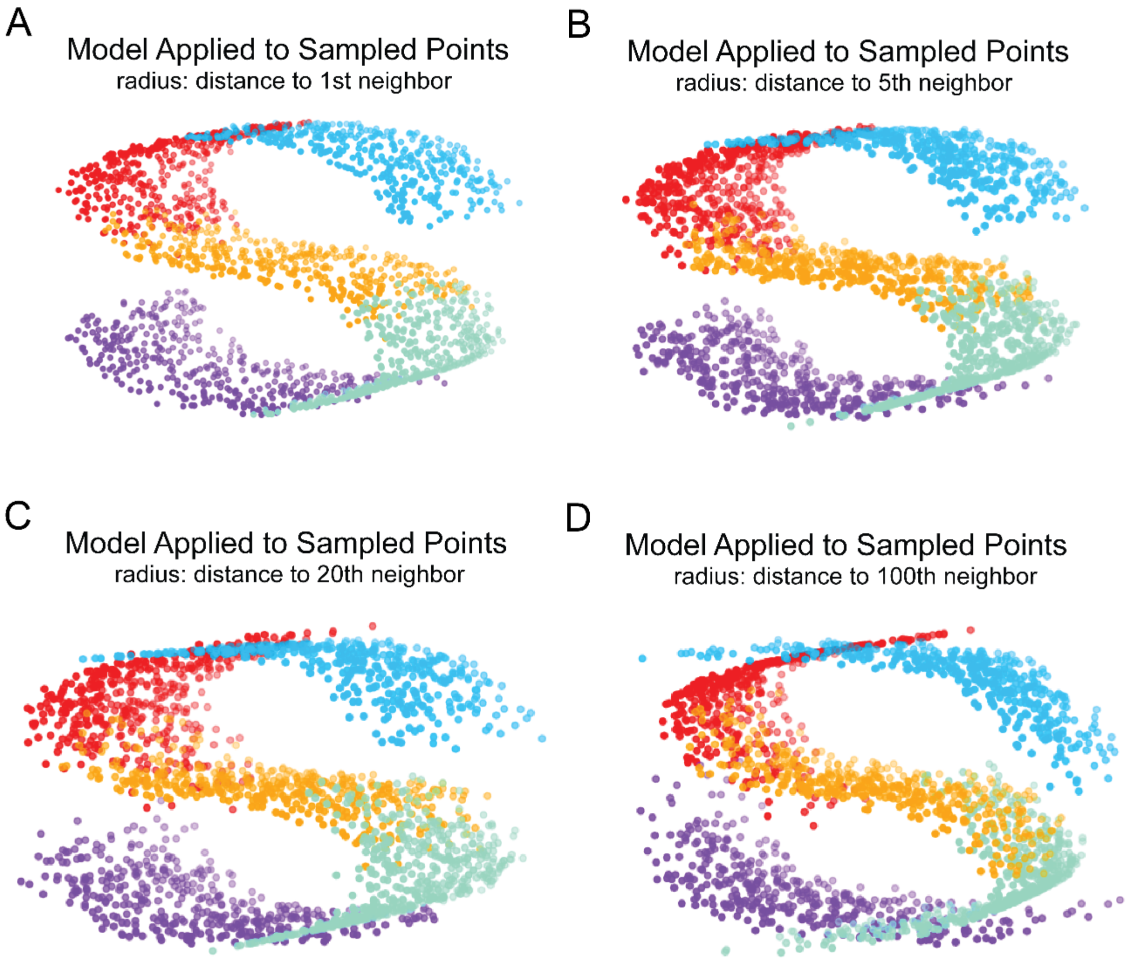


**Fig. S16.**
S-curves generated when applying the model to new data from sampled points. Data generated by sampling from the surface of a sphere with radius of the distance to the 1$^{st}$, 5$^{th}$, 20$^{th}$, and 100$^{th}$ nearest neighbor.
